# High precision *Neisseria gonorrhoeae* variant and antimicrobial resistance calling from metagenomic Nanopore sequencing

**DOI:** 10.1101/2020.02.07.939322

**Authors:** Nicholas D Sanderson, Jeremy Swann, Leanne Barker, James Kavanagh, Sarah Hoosdally, Derrick Crook, the GonFast Investigators Group, Teresa L Street, David W Eyre

**Author notes:** Corresponding author: Nicholas D Sanderson.

## Abstract

The rise of antimicrobial resistant *Neisseria gonorrhoeae* is a significant public health concern. Against this background, rapid culture-independent diagnostics may allow targeted treatment and prevent onward transmission. We have previously shown metagenomic sequencing of urine samples from men with urethral gonorrhoea can recover near-complete *N. gonorrhoeae* genomes. However, disentangling the *N. gonorrhoeae* genome from metagenomic samples and robustly identifying antimicrobial resistance determinants from error-prone Nanopore sequencing is a substantial bioinformatics challenge.

Here we demonstrate an *N. gonorrhoeae* diagnostic workflow for analysis of metagenomic sequencing data obtained from clinical samples using R9.4.1 Nanopore sequencing. We compared results from simulated and clinical infections with data from known reference strains and Illumina sequencing of isolates cultured from the same patients. We evaluated three Nanopore variant callers and developed a random forest classifier to filter called SNPs. Clair was the most suitable variant caller after SNP filtering. A minimum depth of 20x reads was required to confidently identify resistant determinants over the entire genome. Our findings show that metagenomic Nanopore sequencing can provide reliable diagnostic information in *N. gonorrhoeae* infection.

## Introduction

Antimicrobial resistant *Neisseria gonorrhoeae* is a major public health threat, with only limited treatment options available.^1^ We have recently described that rapid long-read sequencing using the Oxford Nanopore Technologies (ONT, Oxford, UK) R9.4.1 platform offers the potential to detect and sequence near-complete *N. gonorrhoeae* genomes directly from urine samples.^2^ This clinical metagenomic approach has the advantage that it does not require prior bacterial culture, which typically adds 2-3 days to diagnostic workflows and may not be available in all cases, particularly in settings where diagnostics are based on molecular testing alone. With analysis possible during sequencing,^3^ it could potentially offer a same day diagnostic tool for gonorrhoea infection that can guide antimicrobial treatment.

ONT data has several potential advantages in addition to speed and the portability of the diagnostic platform. The long reads generated can allow taxonomic classification with greater specificity than is possible with short reads.^4^ Additionally, as reads containing antimicrobial resistance determinants (with the exception of those on plasmids) contain greater amounts of genetic context than is found with short reads, assignment of resistance determinants to a species is more precise. However, ONT data contains a substantial per base error rate of up to 10% with assemblies containing open reading frame disrupting insertion or deletion errors.^5^ Generation of hybrid assemblies with short-read data to mitigate the error rate^6^ negates the speed and portability available with ONT. If Nanopore sequencing is to be used alone for pathogen sequencing applications directly from clinical samples, e.g. for antimicrobial resistance prediction and transmission tracking, then this needs to be overcome.

Previous work,^7^ demonstrates that Nanopore 2D based sequencing of *N. gonorrhoeae* isolates can be used to identify drug resistance determinants and to undertake phylogenetic inference. However, this work was undertaken on isolates, rather than clinical samples directly and Nanopore 2D sequencing has since been deprecated. This study^7^ also found some differences between the phylogenies obtained from ONT and Illumina sequencing of the same isolates as a result of differences in consensus sequences called by the two methods. Most of the previous work optimising consensus sequence calling from Nanopore data has been undertaken following viral sequencing, e.g. of Ebola using Nanopolish.^8^ Some authors have successfully transferred these approaches to bacterial sequence data, e.g. *Escherichia coli* using an optimised application of the GATK package.^9^

Here, we build on this work by releasing a packaged workflow for analysis of 1D R9.4.1 Nanopore data from *N. gonorrhoeae* obtained from direct sequencing of clinical samples. To generate a whole-genome consensus sequence, we use a variant calling approach from aligned reads. For resistance determinant detection we adopt multiple approaches including analysing reads aligned to specific genes. For more diverse genes we use assembled contigs to first select a reference gene before undertaking alignment.

## Methods

We developed an optimised workflow to deliver several outputs from metagenomic sequence data containing *N. gonorrhoeae*: i) classification of sequence reads by species of origin to allow the presence/absence of *N. gonorrhoeae* to be determined, ii) identification of *N. gonorrhoeae* antimicrobial resistance determinants and iii) a consensus whole-genome sequence to facilitate comparisons between genomes for tracking transmission.

### Data sources

To develop and test the performance of our workflow we used ONT data generated in a previous study,^2^ from metagenomic sequencing of *N. gonorrhoeae* nucleic acid amplification test (NAAT)-negative urine samples spiked with varying concentrations of three WHO *N. gonorrhoeae* reference strains: WHO F, WHO V, and WHO X. Additional data from ONT sequences of isolates WHO Q^10,11^ and H18-208^12^ were also used. Details of sequences and accession numbers are given in Table 1. ONT data were compared with Illumina data available for the reference strains and clinical isolates, which was used as a gold standard together with published descriptions of the variants present.^10,12,13^ Illumina data were processed as described previously.^14,15^

**Table 1.**
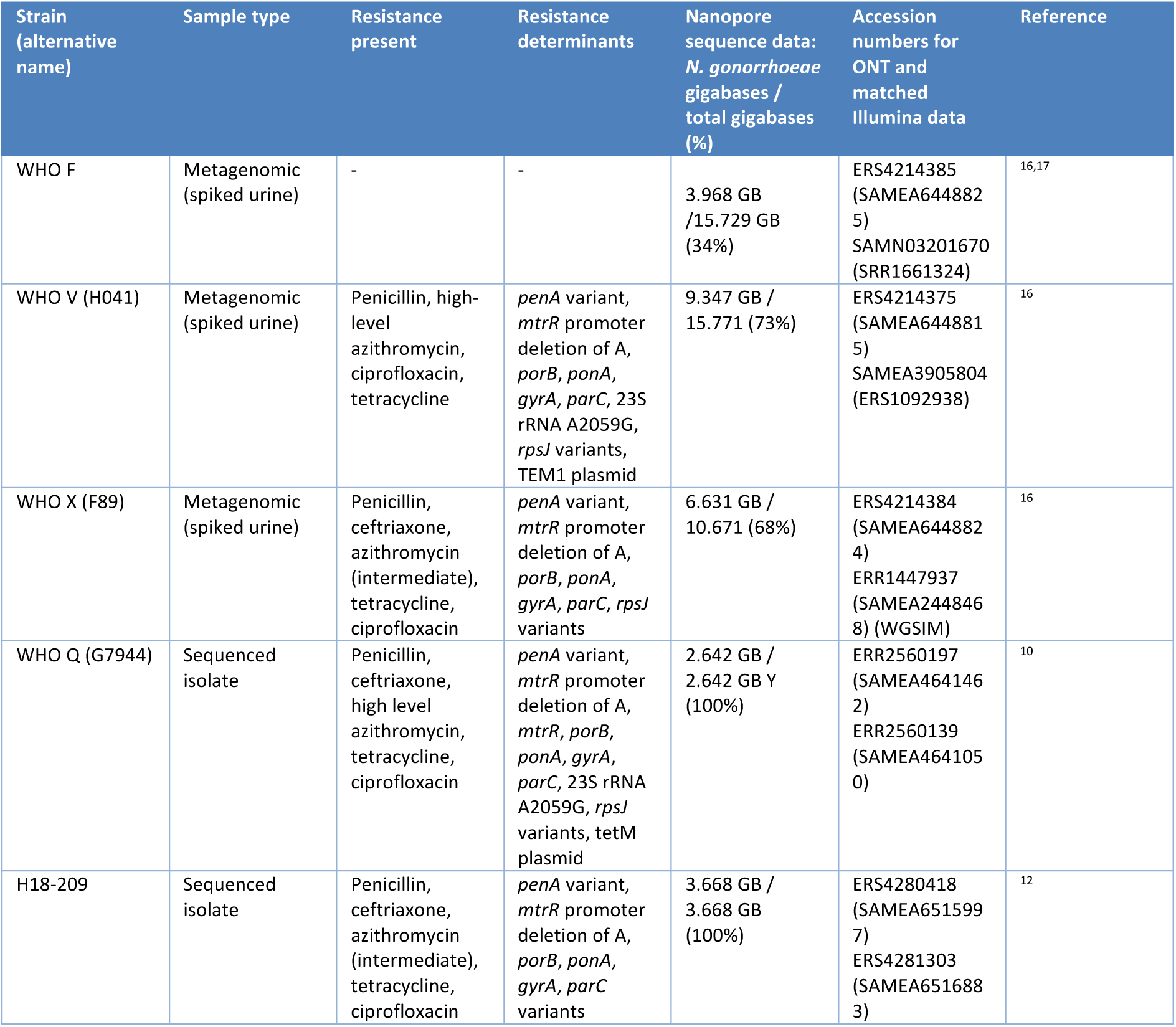
Sequenced isolates and samples. Resistance to azithromycin, ciprofloxacin, tetracycline, penicillin and ceftriaxone is listed with associated genetic determinants. More details on the specific resistance variants can be found in.^10,12,13^

| Strain (alternative name) | Sample type | Resistance present | Resistance determinants | Nanopore sequence data: *N. gonorrhoeae* gigabases / total gigabases (%) | Accession numbers for ONT and matched Illumina data | Reference |
| --- | --- | --- | --- | --- | --- | --- |
| WHO F | Metagenomic (spiked urine) | - | - | 3.968 GB / 15.729 GB (34%) | ERS4214385 (SAMEA6448825)<br>SAMN03201670 (SRR1661324) | 16,17 |
| WHO V (H041) | Metagenomic (spiked urine) | Penicillin, high-level azithromycin, ciprofloxacin, tetracycline | *penA* variant, *mtrR* promoter deletion of A, *porB*, *ponA*, *gyrA*, *parC*, 23S rRNA A2059G, *rpsJ* variants, TEM1 plasmid | 9.347 GB / 15.771 (73%) | ERS4214375 (SAMEA6448815)<br>SAMEA3905804 (ERS1092938) | 16 |
| WHO X (F89) | Metagenomic (spiked urine) | Penicillin, ceftriaxone, azithromycin (intermediate), tetracycline, ciprofloxacin | *penA* variant, *mtrR* promoter deletion of A, *porB*, *ponA*, *gyrA*, *parC*, *rpsJ* variants | 6.631 GB / 10.671 (68%) | ERS4214384 (SAMEA6448824)<br>ERR1447937 (SAMEA2448468) (WGSIM) | 16 |
| WHO Q (G7944) | Sequenced isolate | Penicillin, ceftriaxone, high level azithromycin, tetracycline, ciprofloxacin | *penA* variant, *mtrR* promoter deletion of A, *mtrR*, *porB*, *ponA*, *gyrA*, *parC*, 23S rRNA A2059G, *rpsJ* variants, tetM plasmid | 2.642 GB / 2.642 GB Y (100%) | ERR2560197 (SAMEA4641462)<br>ERR2560139 (SAMEA4641050) | 10 |
| H18-209 | Sequenced isolate | Penicillin, ceftriaxone, azithromycin (intermediate), tetracycline, ciprofloxacin | *penA* variant, *mtrR* promoter deletion of A, *porB*, *ponA*, *gyrA*, *parC* variants | 3.668 GB / 3.668 GB (100%) | ERS4280418 (SAMEA6515997)<br>ERS4281303 (SAMEA6516883) | 12 |

In addition, we also tested our final algorithm on 10 nanopore metagenomic sequences from *N. gonorrhoeae* positive urine samples obtained from men with symptomatic urethral gonorrhoea, described previously.^2^ Cultured isolates from the same infections were sequenced with an Illumina MiniSeq, following the manufacturer’s instructions, to allow for comparisons.

### Workflow

Our end-to-end workflow is written in Nextflow’s domain specific language^18^ and consists of various open source programs and databases (see supplemental material, Figure S1). A second workflow was used to determine the thresholds needed to filter SNPs based on input sequences, truth sequences, and the variant caller used. Both workflows can also be found within a GitLab repository (https://GitLab.com/ModernisingMedicalMicrobiology/ngonpipe).

### Base calling

Raw Nanopore reads were base called with guppy version 3.1.5+781ed57 using the high accuracy HAC models (dna_r9.4.1_450bps_hac.cfg, template_r9.4.1_450bps_hac.jsn). Runs had single barcodes per flow cell so were not demultiplexed.

### Read classification with Centrifuge and read binning

Taxonomic classification of base called Nanopore reads was performed using Centrifuge version 1.0.4-beta^19^, with a database built from NCBI refseq genomes including bacteria and virus genomes deposited as of 10 August 2018 as well as the Human HG18 reference genome. Centrifuge was run with a minimum hit length of 16 (--min-hitlen 16) and reporting a single distinct primary assignment for each read (-k 1). Reads that were classified as, or a strain of, *N. gonorrhoeae* were collected in a separate fastq file using a custom python script (bin_reads.py) available within the GitLab repository.

### Genome alignment

To reduce errors arising from reads from other species mapping to similar genes in the *N. gonorrhoeae* genome, as observed in other metagenomic samples e.g. with *Mycobacterium tuberculosis*,^20^ only reads classified as *N. gonorrhoeae* were aligned. *N. gonorrhoeae r*eads were mapped to the NCCP11945 *N. gonorrhoeae* reference genome (accession NC_011035.1) using minimap2^21^ using settings for Nanopore data (-ax map-ont). Aligned reads were filtered to remove alignments with a map quality score less than 50 and sorted and indexed using samtools.^22^

### Subsampling genome depth

To understand the effect of read depth on variant calling accuracy, aligned bam files for each of the five isolates were subsampled. A custom wrapper script (subSampleBam.py, in the GitLab repository) for “samtools view”^22^ was used to target 2, 5, 10, 20, 50 and 100x average coverage depths.

### Variant calling

Variants were called from the aligned Nanopore reads to either the full genome, or, for variable genes, after remapping to the closest available resistance gene allele from the NG-STAR database (https://ngstar.canada.ca). Several variant callers were tested. Nanopolish version 0.11.1^23^ was used with the methylation aware options (--methylation-aware dcm,dam), --fix-homopolymers, and ploidy set to 1 (--ploidy 1). Medaka version v0.10.0 (https://github.com/nanoporetech/medaka) was used with the consensus and variant subcommands. Clair callVarBam (git commit 54c7dd4)^24^ was used with default ONT settings. Additional information was acquired from pysamstats version 1.1.2 (https://github.com/alimanfoo/pysamstats, pysam 0.15.2) using the variation strand (-t variation_strand) option.

Variants identified by the variant callers were filtered based on metrics generated by pysamstats together with Nanopolish, Medaka, or Clair. Filtering was undertaken using a random forest classifier, using the scikit-learn package,^25^ by comparing Nanopore variant caller outputs and “truth” data from Illumina sequencing of the same isolate. As the purpose of the classifier was to filter potential variants identified by the variant caller, only these sites were used for training. However, summaries of the performance of the classifier at the whole genome level are provided in the results. We defined true positive (TP) SNPs as those that were called and passed by both methods, false positive (FP) Nanopore SNPs that were not found with Illumina sequencing and true negative (TN) sites were called as wild-type by both methods. Sites could be falsely negative (FN) by Nanopore where an Illumina SNP was either missed by the variant caller initially or filtered out incorrectly by the random forest classifier.

To train and test the classifier we used the Nanopore and Illumina sequence for each of the five isolates. To include read depth as a component of the SNP classification, the five genome strains were sub sampled to six target depths of 2, 5, 10, 20, 50, and 100x coverage. All sites from each of the 30 subsampled genomes were randomly divided into a 50% training and 50% validation set. Default hyperparameter values were used. Reported performance metrics include sensitivity or recall, 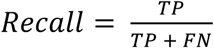 and precision (or positive predictive value for a variant call),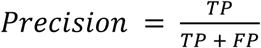.

We considered the following additional metrics obtained using Nanopolish and pysam as input features for the classifier: Variant quality (QUAL), Nanopolish support fraction (Support fraction), total number of reads aligned to each position (Total reads), proximity to the nearest variant in base pairs (proximity), the combination of reference and variant base (baseChange), the proportion of bases the same as the majority base (majority base %), concordance between dominant base and the variant reported (Top base matches variant caller), and proportion of reads that are indels (deletions %, insertions %). This was repeated for Medaka and Clair with the exception of the Support fraction metric that is specific to Nanopolish.

### Indel detection within the *mtrR* promotor

Indels were detected at specific positions within the bam file using a bespoke python script (indel_class.py, available in the GitLab repository), that uses pysam (https://github.com/pysam-developers/pysam) to count the proportion of inserted reads at a position.

### Whole genome assembly (WGA)

Binned reads were filtered for length and quality using Filtlong^26^ for a minimum length of 1000 bp (--min_length 1000), keeping up to 90% of bases (--keep_percent 90) and using a target bases value of 500 mega bases (--target_bases 500000000) as determined in previous work on long read assembly.^6^ Filtered reads were assembled into contigs using Ra^27^ using the -x ont parameters.

### Local gene assembly (LGA) and remapping for *penA* characterisation

For highly variable genes, i.e. *penA*, mapping to a single reference sequence was not possible given the diversity present. Therefore, reads containing genes of interest were identified and isolated using minimap2 and the bin_reads.py script. These local reads were subsequently assembled using wtdbg2 version 2.3^28^ with a longest subread of 3 kb (-L 3000), i.e. the default setting at the time the workflow was developed. A database of available alleles for *penA* was created using the alleles available within the NG-STAR database at https://ngstar.canada.ca. The closest matched allele for each gene was determined using blastn^29^ to search the LGA/WGA contigs. The closest match was chosen as having greater than 95% subject coverage and the highest bitscore. The closest matched allele was then used as a reference to realign binned reads against, using the same mapping and variant calling methods described above.

### *Neisseria gonorrhoeae* antibiotic resistance determinant identification

Following the data processing outlined above, the remaining antimicrobial resistance determinants were identified similarly to our previous approach^15^ developed for short-read sequencing of isolates. Variants in the following genes in the NG-STAR scheme were sought: *penA, mtrR, porB, ponA, gyrA, parC*, 23S rRNA, as well as *rpsJ* mutations and *tet* family genes conferring resistance to tetracycline. Amino acid changes were identified using variant calls in VCF format converted to consensus DNA sequences and then translated. Mutations and variants in promoter sequences were identified from the consensus DNA sequences.

For *penA*, exact matches with one of the alleles in the NG-STAR database were sought (as all isolates / references sequenced were already in the database), but variation from these could also be detected.

To identify mutations in each of the four copies of the 23S rRNA genes associated with macrolide resistance, the four 23S rRNA loci were independently examined for depth of coverage and base changes. This is in contrast to previous approaches using short-read data where the different loci had to be analysed together by mapping to a single copy of the gene.^15^

Antimicrobial resistance conferred by the presence of a specific accessory gene, *e.g.* plasmid associated *tetM/blaTEM-1*, was identified using an assembly strategy. Reads were identified using minimap2 overlaps (-x ava-ont) of all the basecalled reads against a database of accessory gene sequences and assembled with WTBDG2. The resulting contigs were analysed for *tetM/blaTEM-1* sequence and known carrier plasmids for *Neisseria gonorrhoea*e using blastn searches of the same database including pEP5289 (GU479464), pEP5233 (GU479465), pEP5050 (GU479466) for and *tetM*^*30*^ and pEM1 (HM756641.1), pGF1 (U20421), pJD5 (U20375) and pJD7 (U20419) for *blaTEM-1*^*31*^.

### Phylogenetic inference

We compared phylogenetic inferences using Nanopore and Illumina data using whole genome consensus sequences produced after filtering. To reduce the number of false positive and false negative Nanopore SNP calls we also tested additionally masking positions (i.e. setting the base to N) were the proportion of reads supporting the called base was less than a given threshold, e.g. 0.8. Maximum likelihood phylogenetic trees were constructed with IqTree (v1.6.1)^32^ and branch lengths readjusted to account for recombination with ClonalFrameML (v1.11-1)^33^ using default settings. The workflow used is provided within the Nextflow workflow, and is based on runlistcompare (https://github.com/davideyre/runListCompare).

## Results

Data from ONT (Table 1) sequencing of five *Neisseria gonorrhoeae* containing samples were used for initial method development: three metagenomic sequences of urine samples spiked with known reference strains (WHO F, V and X) and two from sequencing of isolates (WHO Q and H18-208). The median sequencing depth was >100x for each sample, and coverage breadth was 97%-99.7% at 1x coverage or higher (Figure S2). Each sequence was subsampled to varying depths between 2x and 100x.

### Tuning variant calling

Variants were called for each subsampled genome using Nanopolish, Clair and Medaka. Previous Illumina sequences of the same isolates were used as a “truth set” or “gold standard” (Table S1). All three variant callers identified numerous false positive SNPs, compared to the Illumina data. Variant caller reported QUAL scores were unable to reliably differentiate false and true SNPs (Figure 1), e.g. using Nanopolish and a QUAL score cut-off of ≥25 for calling variants, at 100x coverage, recall was 0.94-0.97, precision 0.68-0.99, and number of false SNPs 32-1870 across the five genomes. Recall, precision, and false-positive rates for Medaka and Clair were even worse (Figure 1, Table S2).

**Figure 1.**
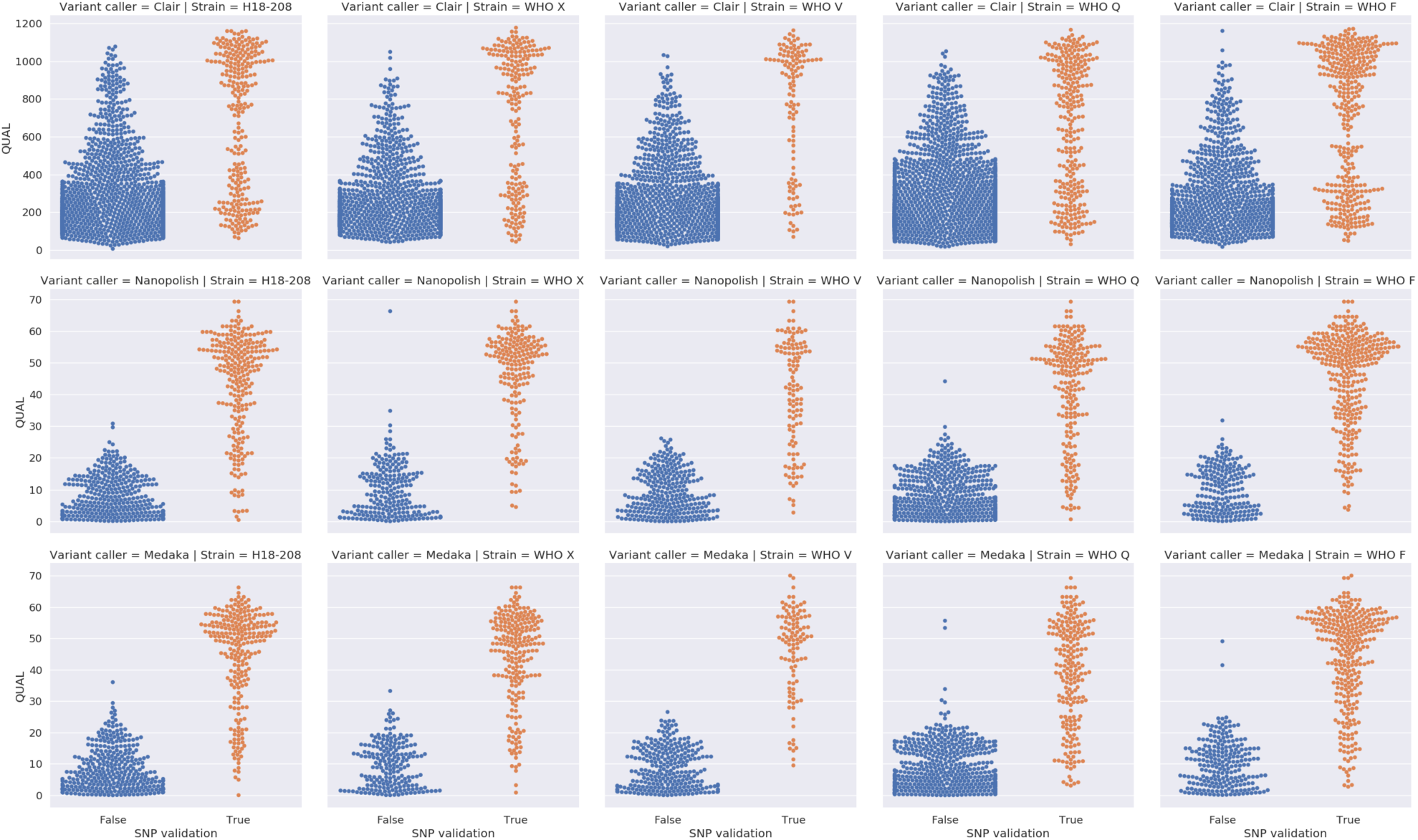
Detection of SNPs using QUAL scores alone. Swarm plots of true (orange) and false SNPs (blue) detected by Clair (top row), Nanopolish (middle row), and Medaka (bottom row). Each column is a different sequence. Each row has different y-axis values.

To improve performance, we trained a random forest classifier to filter the variants using input features from samtools and the variant callers (detailed in the Methods). Performance was assessed using the 50% of bases in the validation set for each genome across all subsampled depths. This approach improved the area under the curve (AUC) for true SNP identification for Nanopolish from 0.86 using a QUAL threshold alone to 0.98 (Figure 2). For Medaka, the AUC improvement was less pronounced, from 0.93 to 0.97. Clair saw the biggest relative improvement from 0.84 to 0.97.

**Figure 2.**
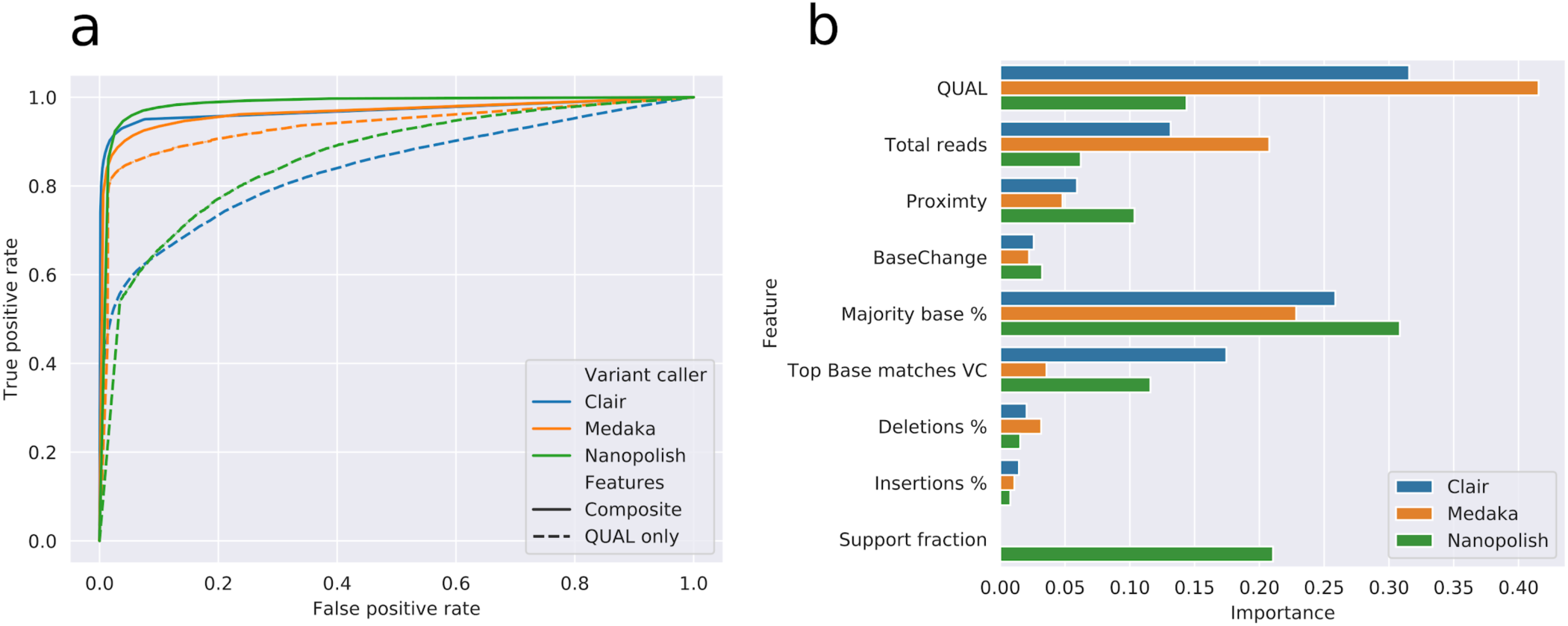
Random forest based variant filtering using Nanopolish, Medaka and Clair. (A) Receiver operator curve (ROC) for random forest classifier using different features including Quality (QUAL only, dashed line) and a composite selection of input features (Composite, solid line) for Nanopolish (green), Medaka (orange) and Clair (blue). Area under the curve (AUC) for each variant caller, Nanopolish 0.86 to 0.98, Medaka 0.93 to 0.97, Clair 0.84 to 0.97, using Qual and Composite features respectively. (B) Bar chart of feature importance for composite selection of features used to train the classifier.

#### Impact of depth of coverage

Using our trained classifier, we assessed the impact of depth of coverage on SNP detection, reporting findings across the whole genome. Increasing coverage up to 20x improved SNP detection, e.g. using Nanopolish, SNP sensitivity was 0.35-0.56, 0.88-0.92 and 0.93-0.95 at 2x, 10x, and 20x coverage respectively across the five genomes (Figure 3A). Medaka had recall rates ∼5% lower than Nanopolish and Clair. Higher coverage depth also reduced the number of false positive SNPs (Figure 3B). Nanopolish had fewest false positives at depths <20x coverage. At 100x coverage, the numbers of false SNPs per genome ranged from 8 to 13 using Nanopolish (i.e. <1 in 100,000 bases), 7 to 28 using Medaka, and 15 to 130 using Clair (Figure 3), with recall rates of 93-95%, 85-92% and 94-98% respectively.

**Figure 3.**
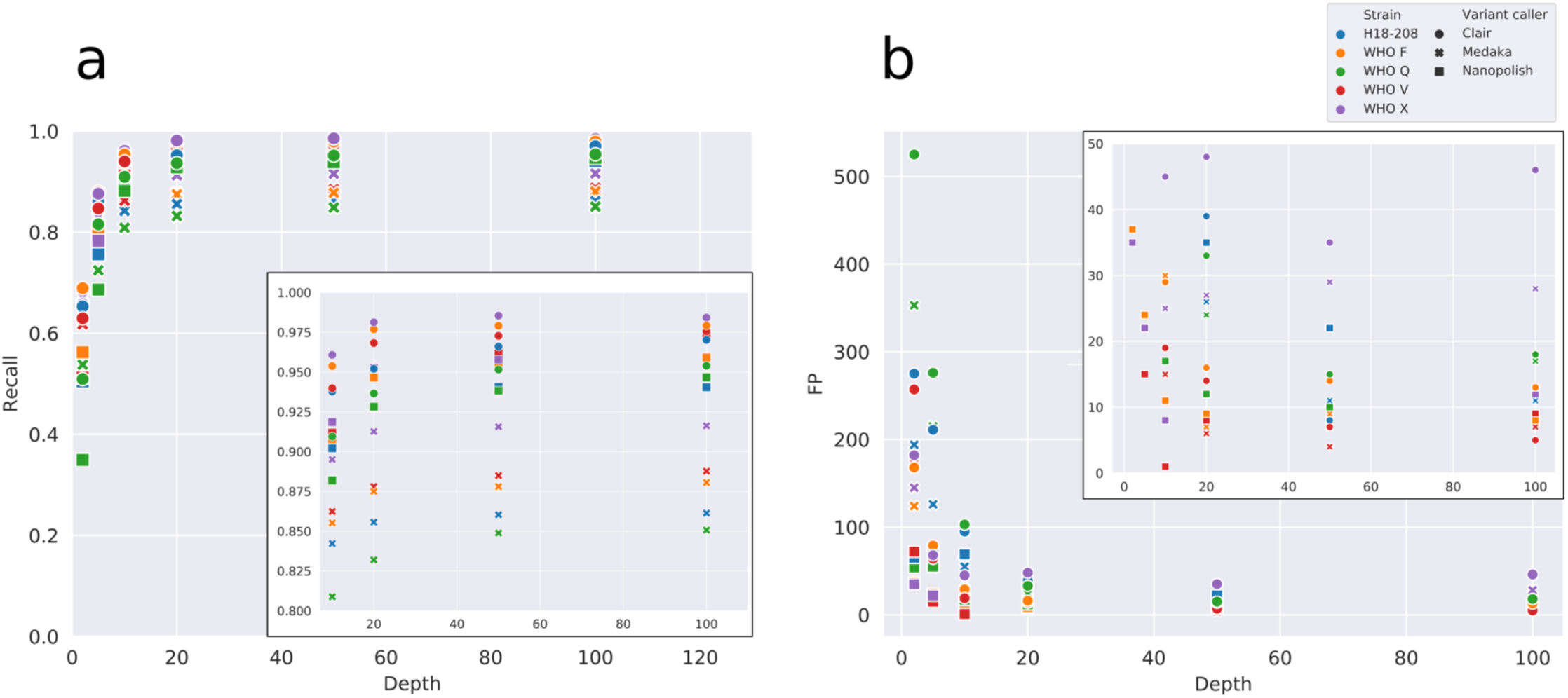
SNP recall by median depth of coverage (A). False Positive SNPs (FP) by median depth of coverage (B). Colour represents different sequences, shapes represent variant callers, circles are Clair, crosses are medaka, squares are Nanopolish. Inserts show upper and lower regions of the y-axis in more detail for panels A and B respectively.

#### Recall performance in important regions and missing SNP calls

We used Clair for subsequent analyses as it offered similar performance to Nanopolish, without requiring resource intensive access to fast5 files. In common with all variant callers tested, Clair missed SNPs (1.5-3%) such that they were not available at the filtering step. If these errors occur systemically, they do not affect comparisons between genomes, however if they occur randomly, they can lead to genomes appearing falsely more similar or different.

Missed SNPs were associated divergence from the reference genome, such that missed SNPs were more closely located to other SNPs (Figure S3). Therefore, for antimicrobial resistance prediction, we only called variants on chromosomal genes with low expected diversity, and selected the closest reference genes for diverse targets, i.e. *penA*. Missed SNPs were not seen within *gyrA, porB, mtrR, parC, ponA* at coverage depths >10x (Figure S4).

### Antimicrobial resistance determinant identification in conserved genes

Antimicrobial resistance determinants were reliably identified by all three variant callers with only a handful of exceptions. All four copies of the 23S rRNA gene were identified separately using long nanopore reads. WHO V and WHO Q contain 4 copies of the A2059G mutation conferring high-level azithromycin resistance. All four mutations were identified at 5x, 10x or 20x coverage using Clair, Nanopolish or Medaka respectively (Table S3). Mutations conferring substitutions at positions 91 and 95 in GyrA and at positions 86-88 in ParC confer ciprofloxacin resistance. These amino acids were correctly identified in GyrA for all genomes at ≥10x coverage with Clair and Nanopolish, but Medaka failed to detect 95N in WHO X at any depth. Expected results were obtained for ParC for all variant callers even at 2x depth (Table S4). Similarly, *ponA* and *rpsJ* mutations (associated with penicillin and tetracycline resistance respectively) were identified at all depths with all variant callers.

Two different types of mutations were examined for the *mtrR* gene, the G45D substitution and promoter variants, which are associated with resistance to azithromycin, ceftriaxone, penicillin and tetracycline. The amino acid at position 45 was corrected called for all genomes at all depths and with all variant callers, except at 2x coverage for WHO Q with medaka (Table S4). A deletion single-base deletion within the promoter, present within all genomes studied except WHO F, was also detected. As the reference sequence contained the deletion, it was expected to be detected as an insertion in WHO F. This insertion was only detected by Nanopolish with 100x coverage. Medaka and Clair detected the insertion at all depths, but also incorrectly identified the insertion in WHO X at ≤5x coverage (Table S5). As indels were not part of our SNP filtering, we developed a heuristic filter for the insertion: 40% or more reads containing an inserted adenosine, with a coverage depth >5x, suggested wild type genotype (Figure S5).

#### penA characterisation using whole genome and local de novo assemblies

The *penA* gene, associated with penicillin and ceftriaxone resistance, is a chromosomal antimicrobial resistance determinant with relatively high nucleotide sequence variation within *N. gonorrhoeae* species arising from recombination events. We identified it using whole-genome and local *de novo* assemblies followed by mapping the closest known allele.

The required average coverage depth to generate contigs containing the *penA* gene was variable between strains (Table S6): H18-208, WHO Q, WHO X, WHO V consistently providing the correct allele with depths of ≥10x. WHO F required 50x coverage for WGA method to recall the allele. The local assembly approach worked for all strains from 10x coverage and higher, it demonstrated better sensitivity at lower read coverage, but did not provide as much genomic context.

#### Detection of plasmid mediated resistance determinants

Plasmid carried *tetM* and *blaTEM-1* confer tetracycline and penicillin resistance respectively. Reads containing *tetM* or *blaTEM-1* sequence were extracted and assembled. To determine if the plasmids were consistent with those in *N. gonorrhoeae* rather than other contaminating species present, we analysed the gene and flanking plasmid sequence. To reliably confirm the presence of these genes contigs containing *blaTEM-1* or *tetM* needed to share >60% sequence proportion matching a known carrier plasmid (Figure S6) with >95% sequence identity. Using this heuristic threshold, it was possible to correctly determine that WHO Q and WHO V contained *tetM* and *blaTEM-1* respectively.

### Longer reads improve metagenomic species disentanglement

To avoid erroneous results arising from DNA from other species only reads classified as *Neisseria gonorrhoeae* to the species level were used for analysis. By limiting the analysis to only this subset of reads, there is a risk of missing regions of the genome by filtering reads that assign to a lower taxon^34^. We therefore tested the expected proportion of the *N. gonorrhoeae* genome that would be classified to the species level by simulation (Figure S7). In contrast to other species, *N. gonorrhoeae* could reliably be identified to the species level with read lengths of a few hundred base pairs. The mean read length from our sequencing was between 2-4 kb^2^, which enabled a high proportion of *N. gonorrhoeae* sequence to be assigned to the species level.

### Further filtering to remove false SNP calls

When using SNP data to reconstruct transmission events, false SNPs can lead to transmission being incorrectly excluded or deemed unlikely. Similarly, missed SNPs occurring at random, where the consensus sequence is wrongly set to be wild-type can increase measured genetic distance between two similar strains. In contrast, the expected sequence difference at filtered sites, where the base is unknown, can be adjusted for in proportion to the percentage of the genome filtered and variation in the known genome. Therefore, for transmission studies, a strategy of favouring removing false-positive and false-negative SNPs over recall is preferred. To achieve this, the SNP classifications were further filtered by masking nucleotide classifications to N if the proportion of bases at a given position supporting the classification was <0.8. This value was chosen as the proportion of true-positive SNPs with support <0.8 is relatively low, but this threshold is sufficiently high to avoid most false negative calls (Figure S8).

By using this final filter with Clair base-called data at 100x coverage, the number of false-positive SNPs was reduced from 15-130 to 9-35 across the five genomes analysed, Table 2. The number of false-negative SNPs also fell from 49-249 to 4-19. Overall this resulted in false SNP rates (false negative + false positive SNPs) falling from 66-428 to 15-45, with a reduction in recall from 0.93-0.99 to 0.76-0.94, which is likely to still remain acceptable for most transmission studies.

**Table 2.** Recall rates for filtered and unfiltered spiked genomes, variant called with Clair and a random forest classifier. Show with (“Filtered) and without (“Unfiltered”) additional filtering by requiring 80% of bases to support the called nucleotide. Data shown for 100x coverage.

| Sample | Total SNP positions | Filtered FN | Filtered FP | Filtered TP | Unfiltered FN | Unfiltered FP | Unfiltered TP | Unfiltered Recall | Filtered Recall |
| --- | --- | --- | --- | --- | --- | --- | --- | --- | --- |
| H18-208 | 4297 | 12 | 9 | 3740 | 289 | 130 | 4008 | 0.93 | 0.87 |
| WHO F | 6095 | 19 | 12 | 5660 | 130 | 15 | 5965 | 0.98 | 0.93 |
| WHO Q | 3844 | 12 | 4 | 2913 | 167 | 16 | 3677 | 0.96 | 0.76 |
| WHO V | 1799 | 4 | 11 | 1620 | 49 | 17 | 1750 | 0.97 | 0.90 |
| WHO X | 3366 | 10 | 35 | 3153 | 50 | 45 | 3316 | 0.99 | 0.94 |

### Application of the workflow on clinical samples

We analysed previously generated nanopore metagenomic sequencing data from ten urine samples from men with urethral gonorrhoea. We compared findings with our workflow to Illumina data obtained as part of this study from sequencing isolates from the same infections. By nanopore sequencing, ≥92.8% coverage of an *N. gonorrhoeae* reference genome was achieved in all samples, with ≥93.8% coverage breath at ≥10-fold depth in 7.

All resistance gene SNPs were correctly identified in the metagenomic clinical samples (Table S7). Using the heuristic method, the *mtrR* promoter deletion was correctly detected in samples 202, 250, 301 and 314, and the wild-type sequencing in samples 271, 294 and 315. However, sample 303 was incorrectly identified, with only 11x mean genome coverage depth and 8x coverage over the *mtrR* gene suggesting a lack of sequencing depth to accurately call the position, as in Table S7. The *penA* allele was correctly identified in 9 of the 10 clinical samples (Table S8). All clinical metagenomic samples identified corresponded with Illumina sequenced cultures at 100% identity according to blastn results. Sample 303 produced insufficient data to detect the *penA* gene. It was also possible to determine that samples 206, 271, 294 and 304 contained the *tetM* gene on the pEP5050 plasmid, and samples 294 and 303 contained the *blaTEM-1* gene on the pEM1 plasmid (Figure S8).

By producing a Nanopore consensus sequence with only high probability SNPs added, and sites with <80% support set to N, i.e. unknown, conventional tree building methods can be used. This approach demonstrated comparable findings between cultured isolates sequenced with Illumina and clinical metagenomic samples sequenced with Nanopore (Figure 4). Samples 303 and 304 provided insufficient data to generate complete consensus sequences (only 53 and 56% of the reference genome length was identified). For the remaining 8 clinical samples and 5 method development sequences the median (IQR) [range] genetic distance between the Illumina and Nanopore sequences from the same infection was 5 (3-6) [1-10] SNPs, which is close enough to make transmission studies possible using metagenomic data alone.

**Figure 4.**
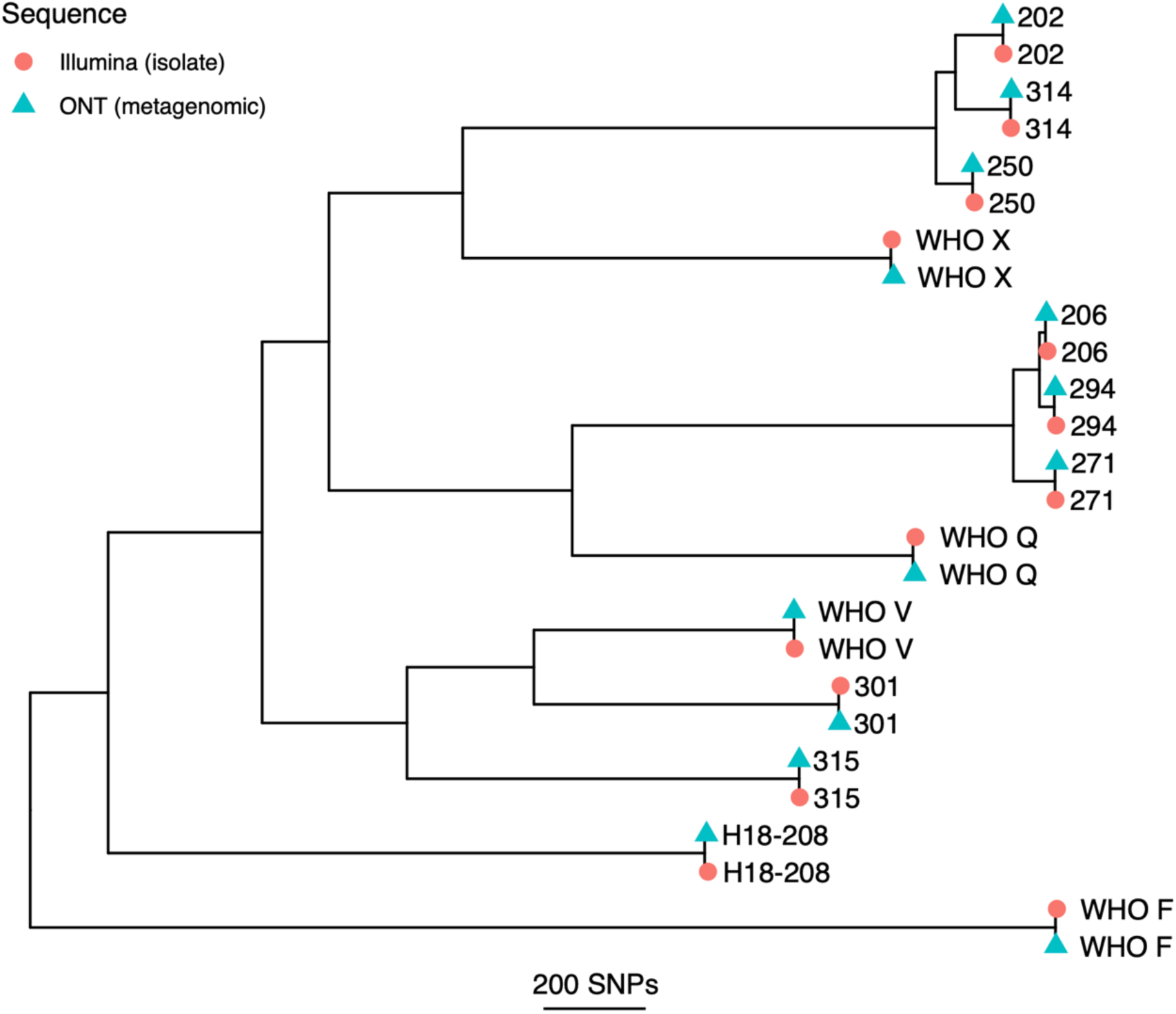
Recombination-corrected maximum likelihood tree of metagenomic nanopore and paired Illumina isolate sequences. All nanopore consensus sequences were generated from metagenomic sequencing with the exception of H18-208 and WHO Q which were sequenced from isolates.

## Discussion

We demonstrate an approach that allows Nanopore sequencing data to be used to reconstruct accurate consensus bacterial genomes. This can be done without accompanying Illumina short-read data and can be applied to metagenomic sequencing data. We show the reconstructed genomes allow accurate resistance detection and transmission inferences to be made in *N. gonorrhoeae*, including using samples obtained from clinical infections.

We evaluated three variant callers, Nanopolish, Medeka, and Clair, against Illumina variant calling from sequenced cultures. After filtering variant calls with a trained random forest classifier, we found that Clair performed better than Nanopolish and Medaka, identifying 94-98% of SNPs present in Illumina sequences at 100x coverage, compared to 93-95% and 85-92% respectively. However initially Clair had the highest number of false positive SNPs per genome (15-130, compared to 8-13 and 7-28 respectively). By using further filtering, requiring the proportion of reads supporting any call to be ≥0.8, the number of false positive SNPs could be reduced using Clair to 4-35/genome, albeit with a reduction in SNP detection to 76-94%.

Importantly, this filtering and masking approach also reduced the number of false-negative SNPs from 49-289/genome to 4-19/genome, that would otherwise increase genetic distance during phylogenetic inference. For variant detection in resistance genes specifically, Clair was able to detect all the important SNPs with a coverage of 10x and above, whereas Medaka missed an important SNP in the WHO X strain.

Medaka (v0.10) is still an early release experimental research tool that is focused more on diploid variant calling and haplotype phasing rather than the application tested here. Medaka and Clair have the advantage of not requiring the fast5 files which have a huge storage and computational requirement. The threshold analysis workflow written here has been designed to drop in different variant calling components, to allow for testing new variant calling options in future.

As our Illumina data truth set pipeline only produced SNP calls, our current variant call filtering was limited to SNPs. Indels were not considered except for the *mtrR* promoter region where a bespoke heuristic method was used. Therefore, further work on the Illumina sequences will be needed to provide a truth set for indel data to allow the development of robust indel calling from Nanopore data, which may also improve with future Nanopore pore technology.

By subsampling reads to produce artificially reduced coverage depths, we have determined the required depth needed to accurately call variants from Nanopore data: 10x fold coverage is sufficient to define resistance determinants with minimal increase in recall above 20x fold coverage.

We were successfully able to detect relevant *N. gonorrhoeae* antimicrobial resistance determinants conferring resistance to clinically important antibiotics across all samples tested with a coverage depth above 10x. Most variants could be detected from appropriately filtered variant calls from mapped data and *penA* allele determination could be achieved using a combined assembly and mapping approach. The WGA approach provided more genomic context around the *penA* allele, that could guarantee the allele was from *Neisseria gonorrhoeae* and not a contaminating commensal, whereas LGA performed better at lower read depths. Ra was chosen as the assembler for WGA as WTDBG2 (redbean) produced some mis-assembly that prevented remapping of reads to the *penA* locus (data not shown). However, Ra failed to produce contigs for most of the attempts when used for the LGA. It was possible to recover the four 23S rRNA loci separately from each sample containing the expected A2059G mutation. This was not possible using short read Illumina sequencing. The Nanopore read length allowed us to span the entire gene with enough genomic context to confidently map to each locus independently.

Our final consensus sequence comparison yielded a median of 5 SNPs between Illumina and Nanopore sequences. Although this does not match the reproducibility seen in Illumina sequencing of isolates,^14^ it is close enough to judge whether infections are part of specific transmission clusters, even if precise reconstruction of individual transmission events may remain challenging with Nanopore data alone.

The current generation of ONT flow cells used in this analysis is R9.4.1. However, new pores such as R10 are currently in development and may offer increased accuracy. The validation part of this workflow should be run on new sequences generated by future pores to set new threshold values and filtering models that are appropriate to these new pore error profiles.

The approaches we have developed provide a mechanism for determining antimicrobial resistance and undertaking transmission tracking using clinical samples. This, taken together with recent advances in optimising DNA extraction for metagenomic Nanopore sequencing of *N. gonorrhoeae* direct from urine samples^2^, this now provides an opportunity to test the performance of Nanopore sequencing as a clinical diagnostic in *N. gonorrhoeae* infection. Furthermore, this approach may have wide applicability across a range of bacterial pathogens, not just *N. gonorrhoeae*. Evaluations in clinical datasets will allow the potential utility of our approaches to be further investigated and potentially provide new diagnostics to guide patient and public health management of gonorrhoea.

## Data availability

Nanopore data is available in project accessions PRJEB35173 and PRJEB26560. Illumina sequenced culture isolates are available in project accession PRJNA603903. Data analysis workflow is available as git repository (https://gitlab.com/ModernisingMedicalMicrobiology/ngonpipe.)

## Acknowledgements

The authors thank the microbiology laboratory staff of Oxford University Hospitals NHS Foundation Trust and the Royal Sussex County Hospital, Brighton for providing assistance with sample collection.

The GonFast Investigators Group includes Joanna Rees and Emily Lord (Oxfordshire Sexual Health Service, Oxford University Hospitals NHS Foundation Trust, Oxford, UK) and Suneeta Soni, Celia Richardson, Joanne Jessop, and Tanya Adams (Brighton and Hove sexual health and contraception service, Royal Sussex County Hospital, Brighton, UK), and Martin Llewelyn (Royal Sussex County Hospital, Brighton, UK).

## Conflicts of interest

DWE has received lecture fees and expenses from Gilead unrelated to the current study. No other author has a conflict of interest to declare.

## Funding

This work was funded by the Centers for Disease Control and Prevention, under Broad Agency Award FY2018-OADS-01.

## Supplemental Figures

**Supplemental Figure S1.**
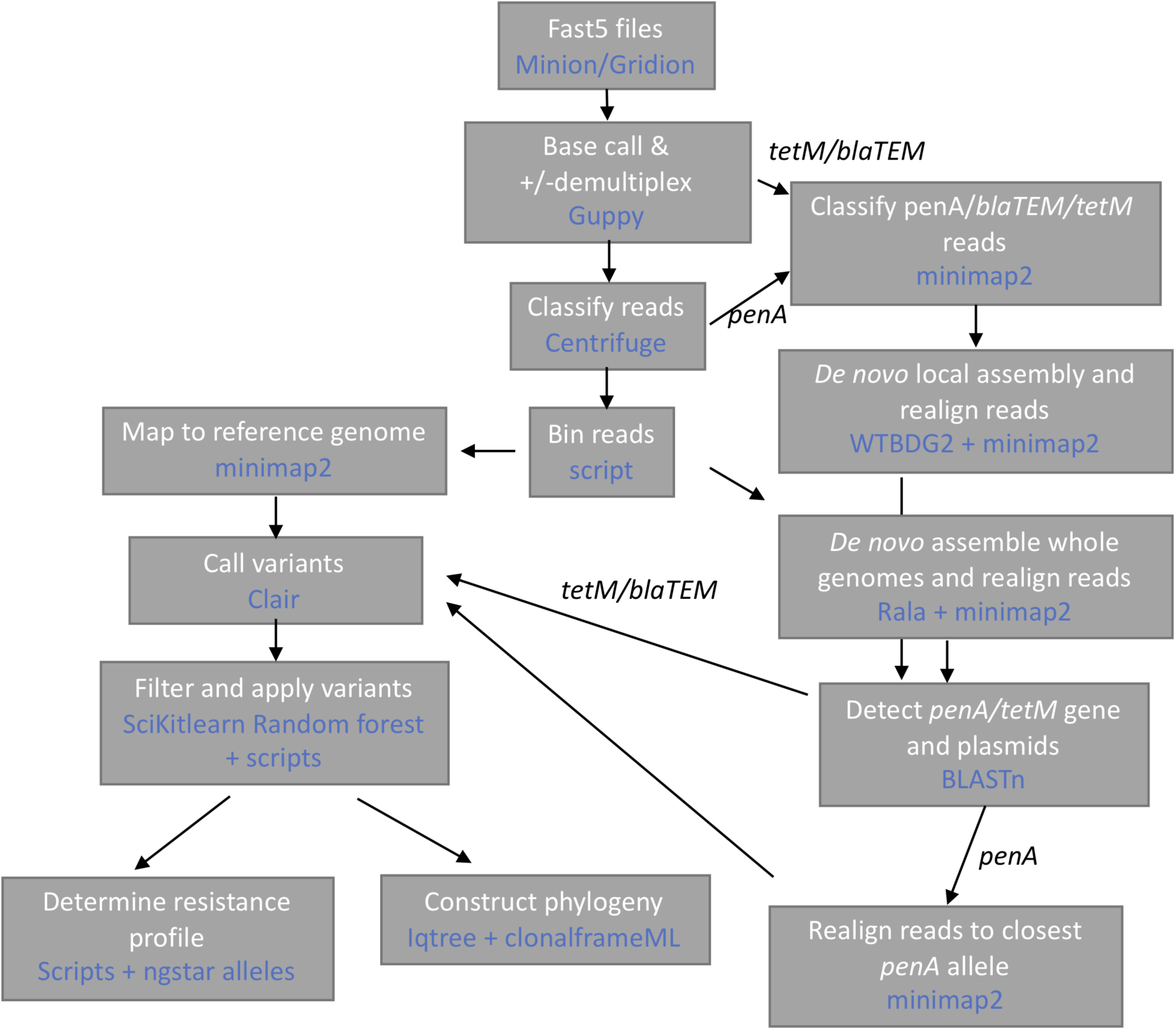
Basic workflow schematic. For antimicrobial resistance prediction mtrR, porB, ponA, gryA, parC, 23S rRNA are identified from the consensus sequence obtained from mapping to the default reference, penA uses a custom reference -see right-hand branch of schematic. Genes present on plasmids (tetM, blaTEM) are identified by mapping to a collection of accessory genes.

**Supplemental Figure S2.**
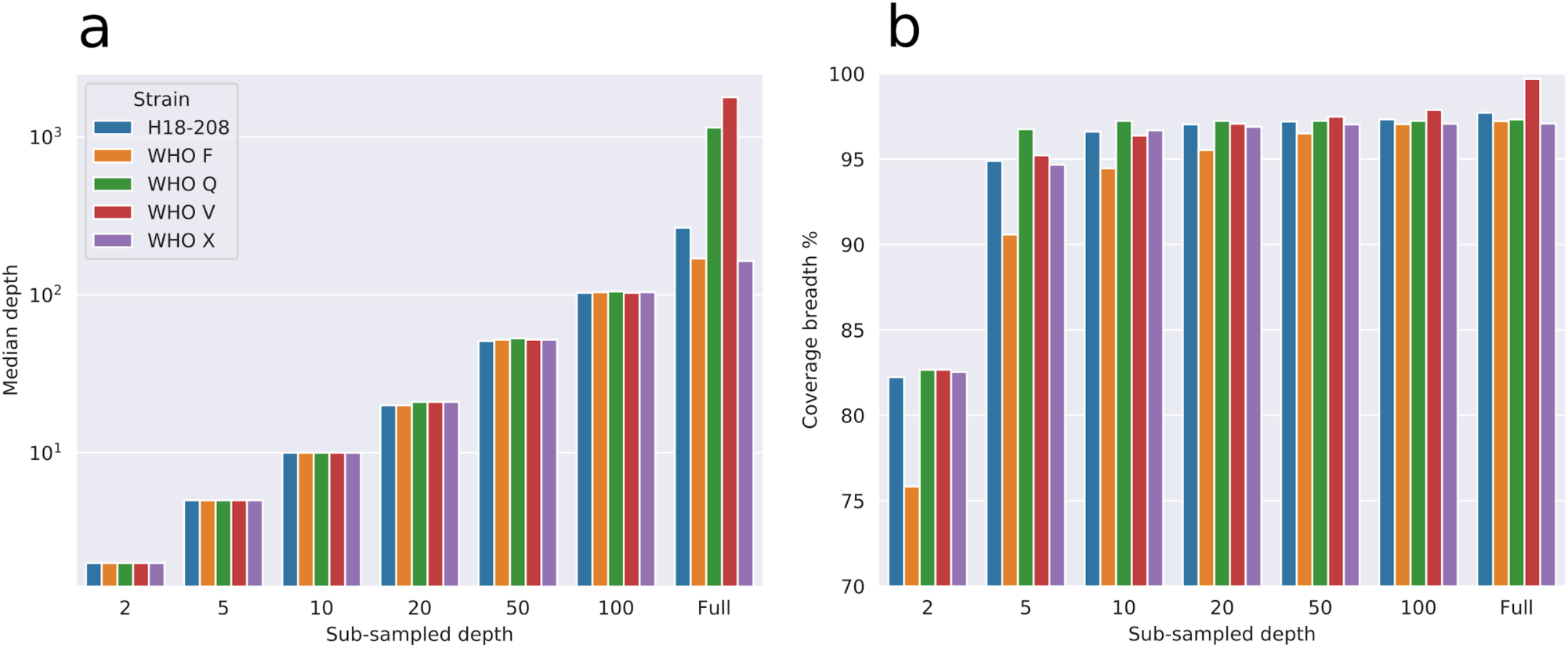
Sequence coverage depth and breadth in original and subsampled data. (A) Median depth achieved from subsampling bam files and original unsampled bam files (Full). (B) Coverage breadth at each target depth.

**Supplemental Figure S3.**
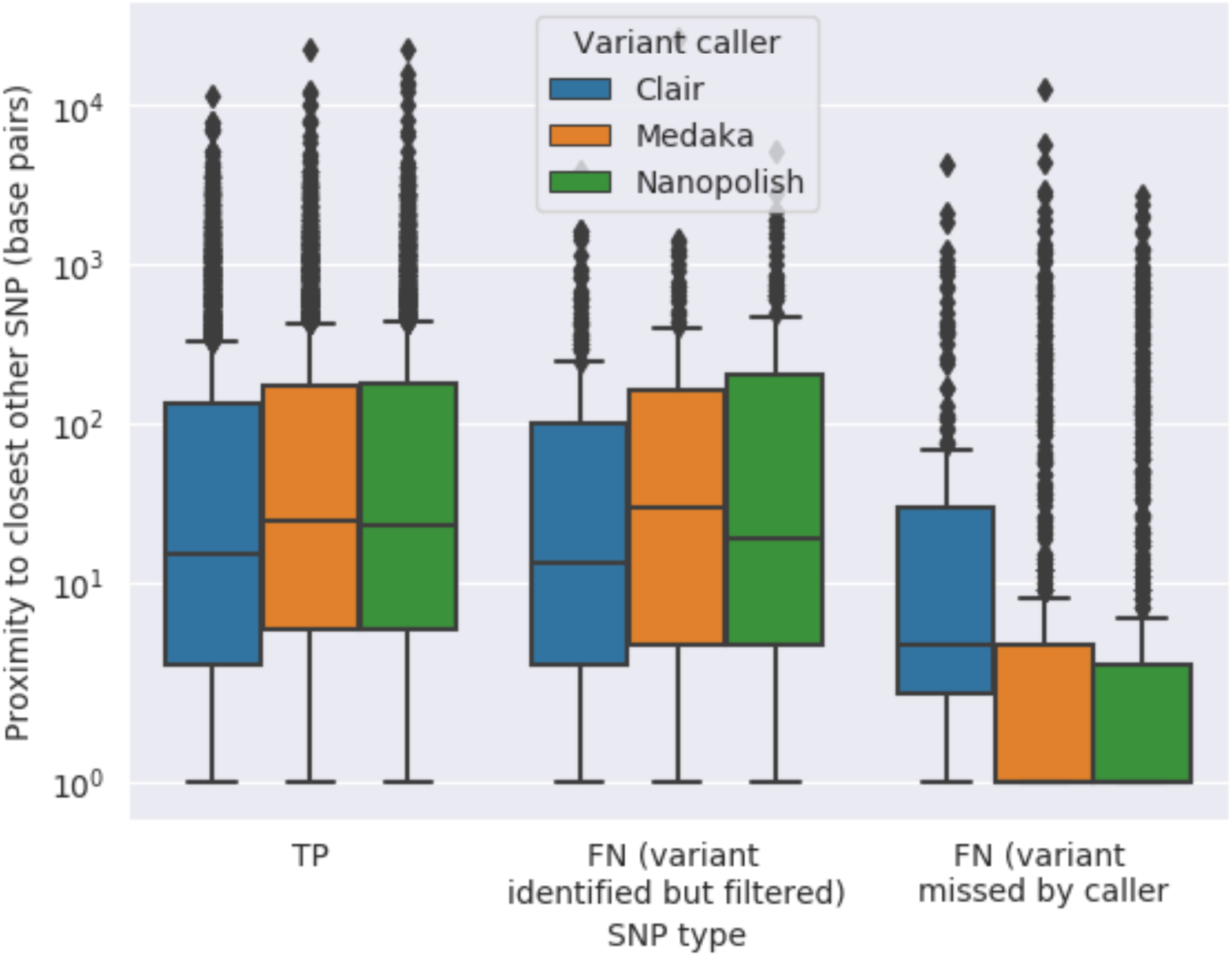
box and whisker plots of detected and missed SNPs and proximity to closest genuine SNP using each variant caller.

**Supplemental Figure S4.**
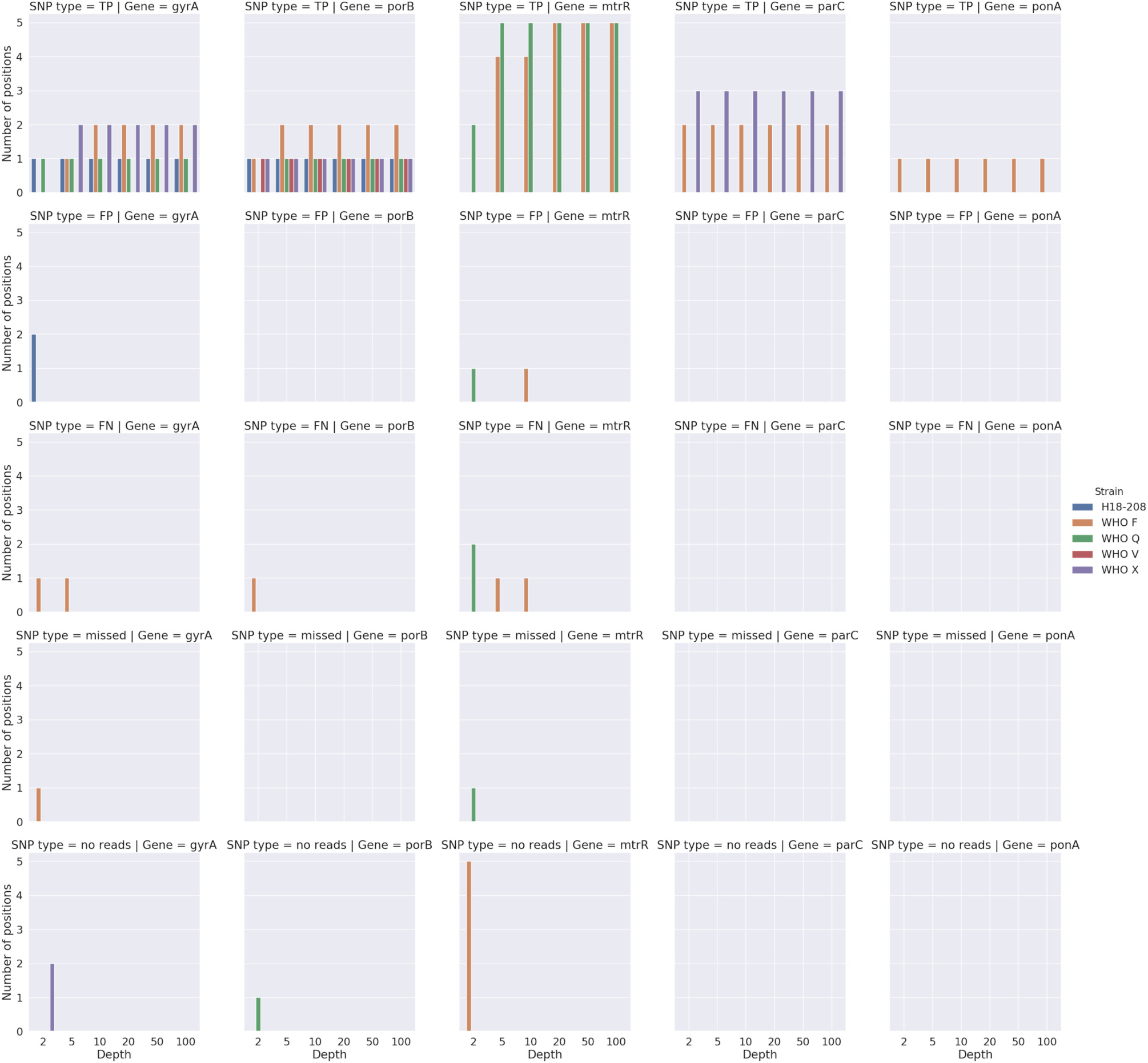
SNP detection in N. gonorrhoeae genes associated with antimicrobial resistance. Bar plots showing number of positions with different SNP types (columns) detected by Clair in important genes (rows). From left to right: gyrA, porB, mtrR, parC, ponA. From top to bottom: True positives (TP), False positives (FP), False negatives (FN), Missed SNPs, no reads covering gene. All plots share a y-axis axis of number of positions classified and x-axis of target depth of coverage. Colours represent each sequence used. The 23S gene was masked from Illumina mapped reads as the read lengths were too short to distinguish different loci and are therefore not compared here.

**Supplemental Figure S5.**
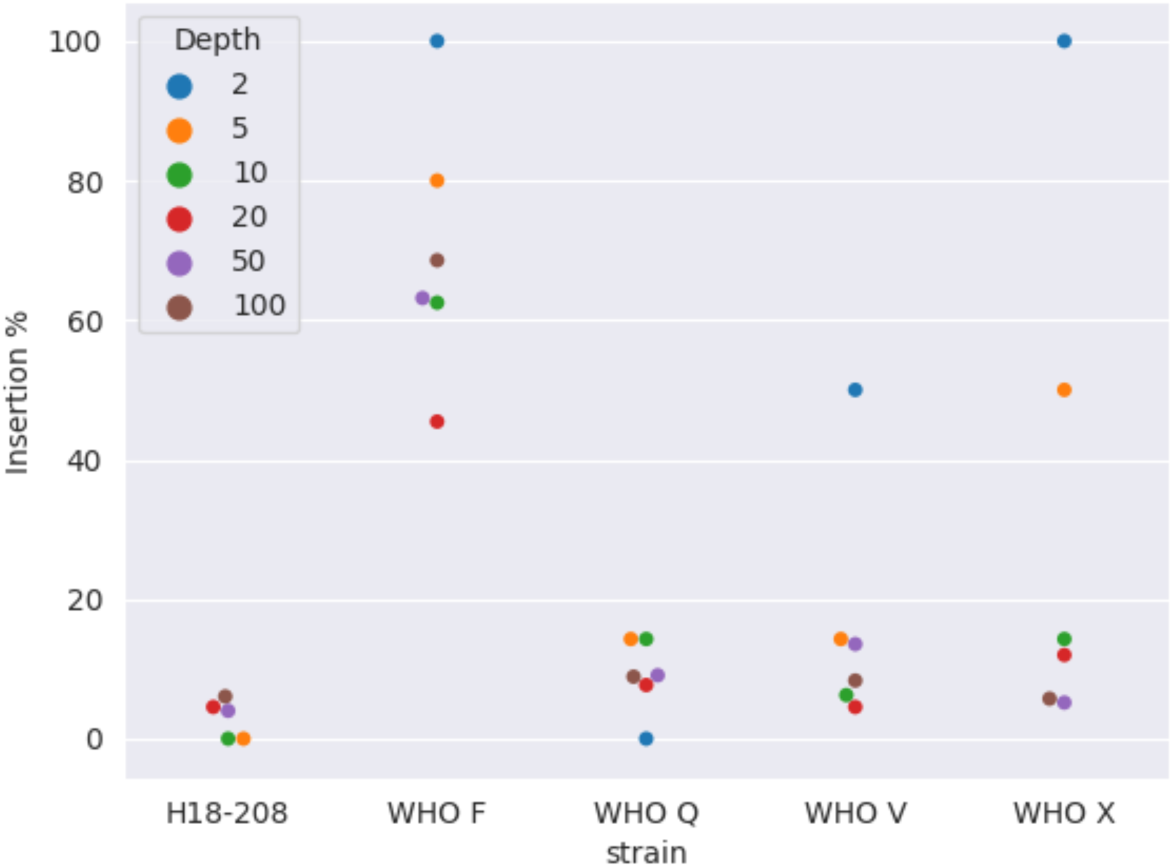
mtrR promoter deletion of adenosine base. The reference genome used contains the deletion, and the true wild-type is represented by an insertion. Percentage of reads with an inserted base at position 1332810 in reference NC_011035.1.

**Supplemental figure S6.**
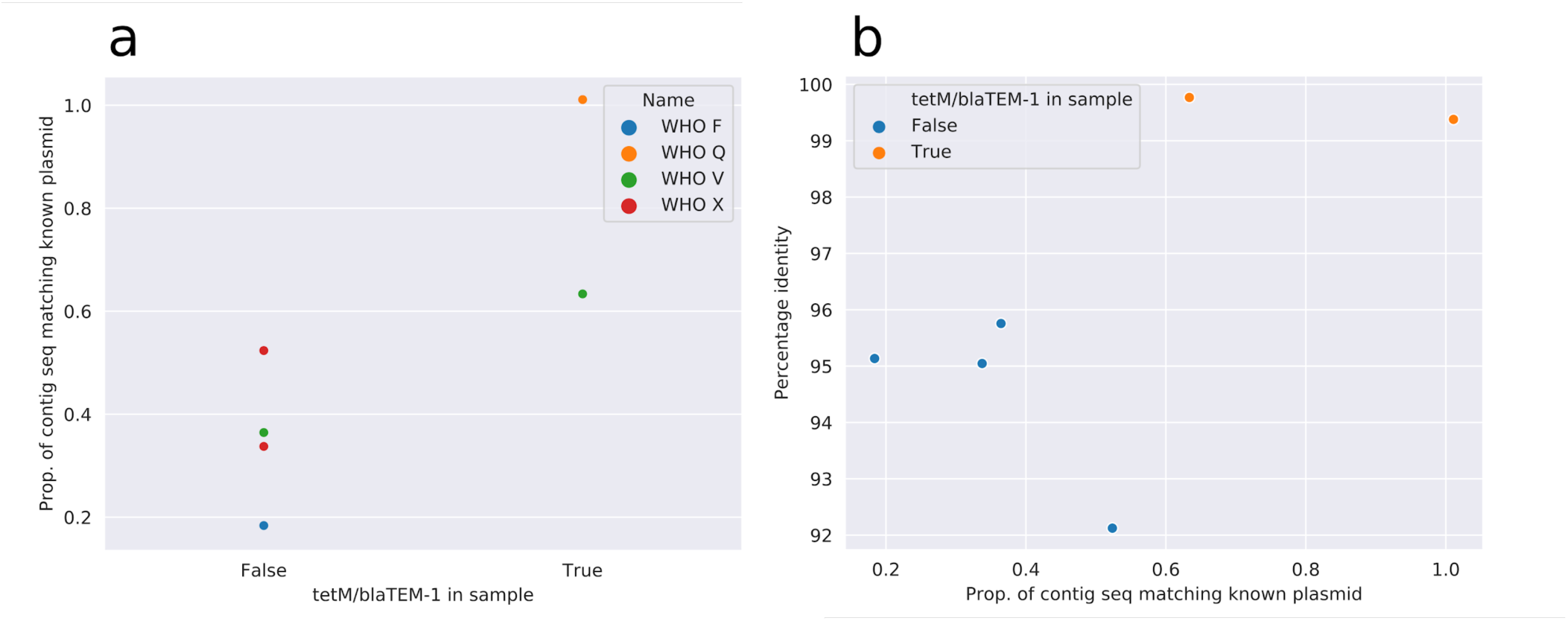
detection of plasmid-borne resistance genes. Proportion of the query sequence (assembled contig) matching a known carrier plasmid (y-axis) over validation of the sample containing tetM or blaTEM-1 (x-axis), colours represent samples (A). Percentage identity between the matching contigs and carrier plasmid (y-axis) over proportion of the query sequence matching a known carrier plasmid (x-axis), colour represents validation of containing tetM or blaTEM-1 (B).

**Supplemental figure S7.**
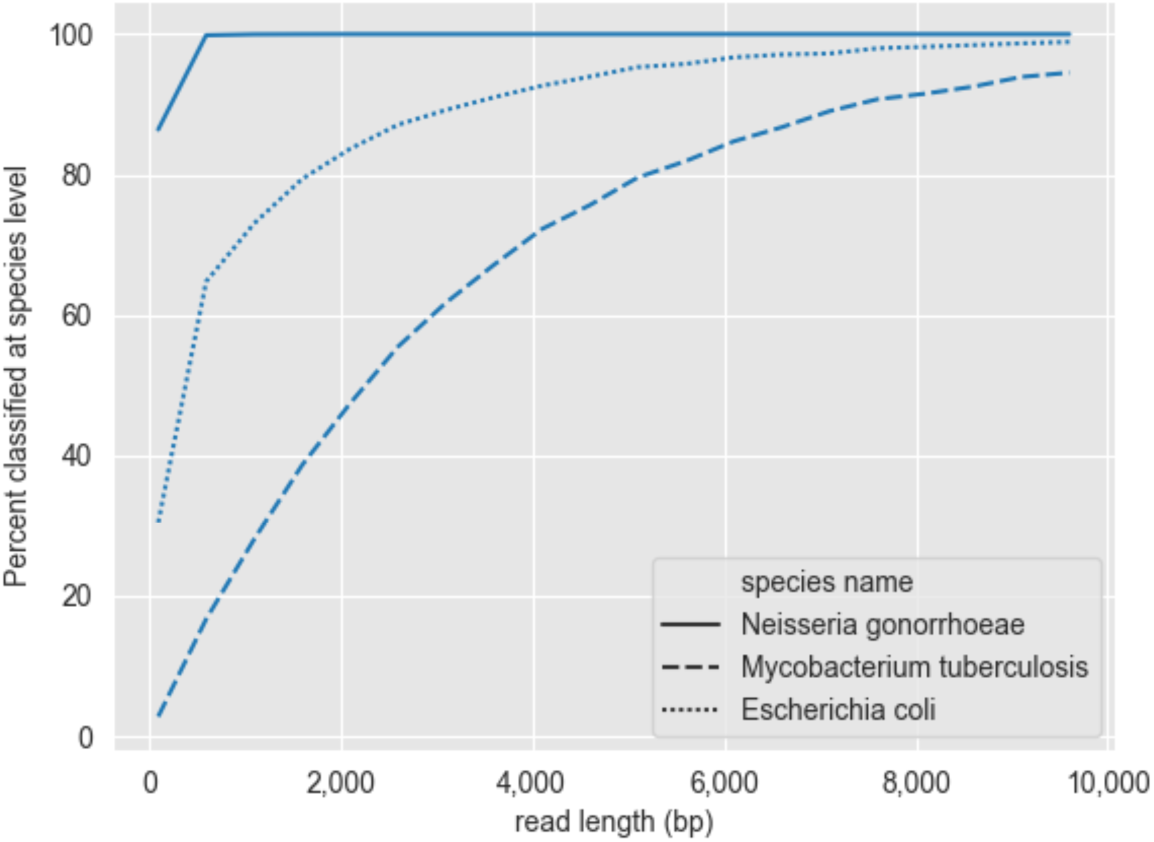
Taxonomic assignment by read length. Simulated raw reads for N. gonorrhoeae and two comparison species.

**Supplemental Figure S8.**
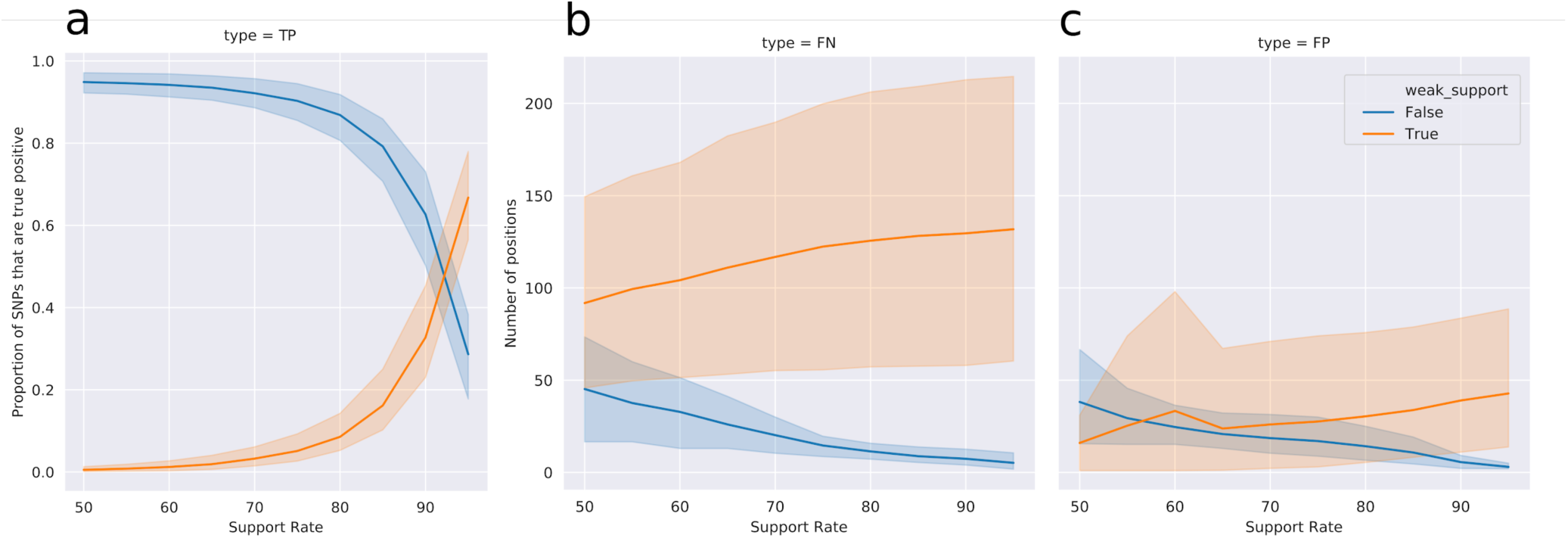
Filtering SNP calls based on support rate. Proportion of SNPs that are true positives classified by weak support (True equates to less than the support rate, orange or False equates to more that the support rate, blue) over support rate (A). The number of positions that are False Negatives (B) or False Positives (C) over support rate and classified by weak support (same as A).

**Supplemental figure S9.**
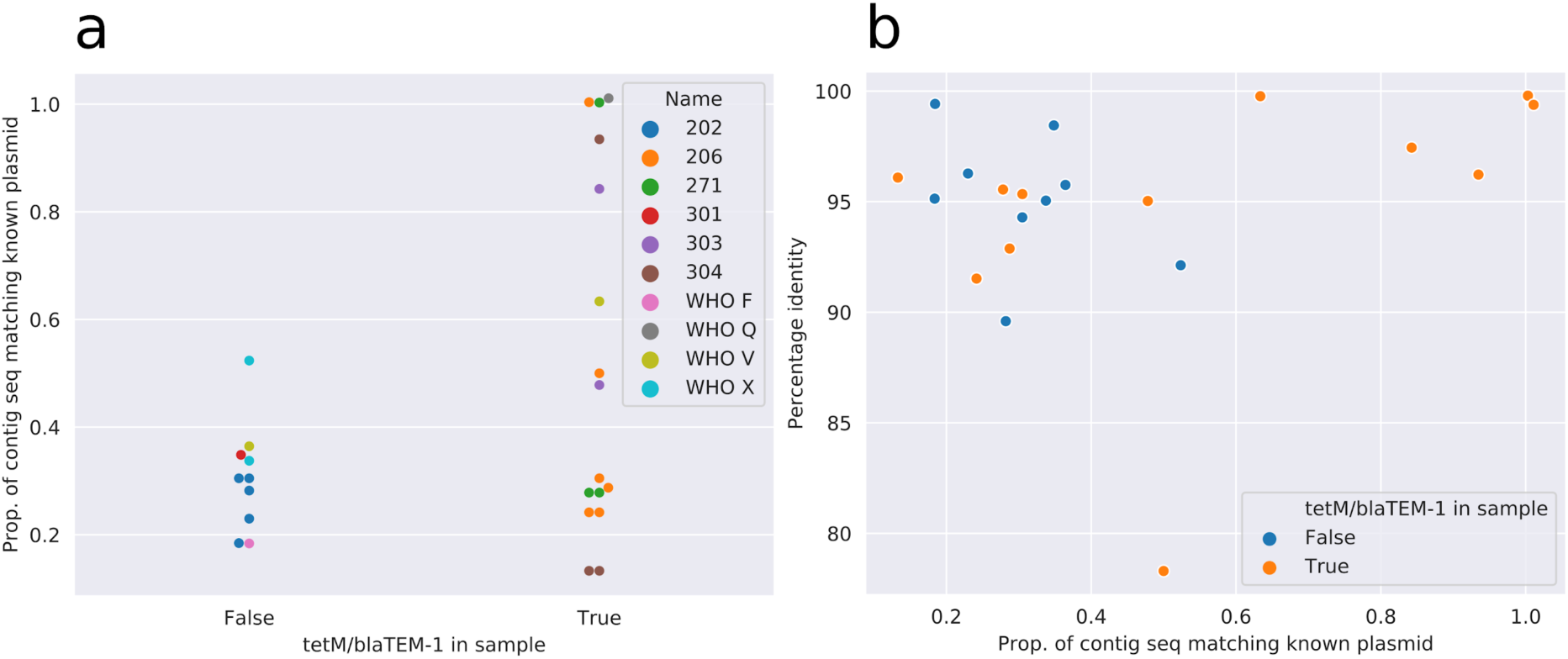
Plasmid-borne resistance gene detection in clinical samples. Proportion of the query sequence (assembled contig) matching a known carrier plasmid (y-axis) by presence of tetM or blaTEM-1 in the sample (x-axis), colours represent samples (A). Percentage identity between the matching contigs and carrier plasmid (y-axis) over proportion of the query sequence matching a known carrier plasmid (x-axis), colour represents validation of containing tetM or blaTEM-1 (B).

## Supplemental Tables

**Supplemental Table 1.** “Gold standard” variation present from Illumina sequencing. Number of variants, invariant sites, and uncalled sites found within each sample from Illumina sequencing.

| Strain | Illumina bases covered | Reference Chromosome length | Missed (N) sites | Illumina SNPs | Invariant sites |
| --- | --- | --- | --- | --- | --- |
| H18-208 | 1,992,319 | 2,232,025 | 239,706 | 4,303 | 1,988,016 |
| WHO Q | 1,962,756 | 2,232,025 | 269,269 | 3,844 | 1,958,912 |
| WHO F | 1,991,234 | 2,232,025 | 240,791 | 6,174 | 1,985,060 |
| WHO V | 2,016,466 | 2,232,025 | 215,559 | 1,801 | 2,014,665 |
| WHO X | 1,955,978 | 2,232,025 | 276,047 | 3,366 | 1,952,612 |

**Supplemental Table 2.**
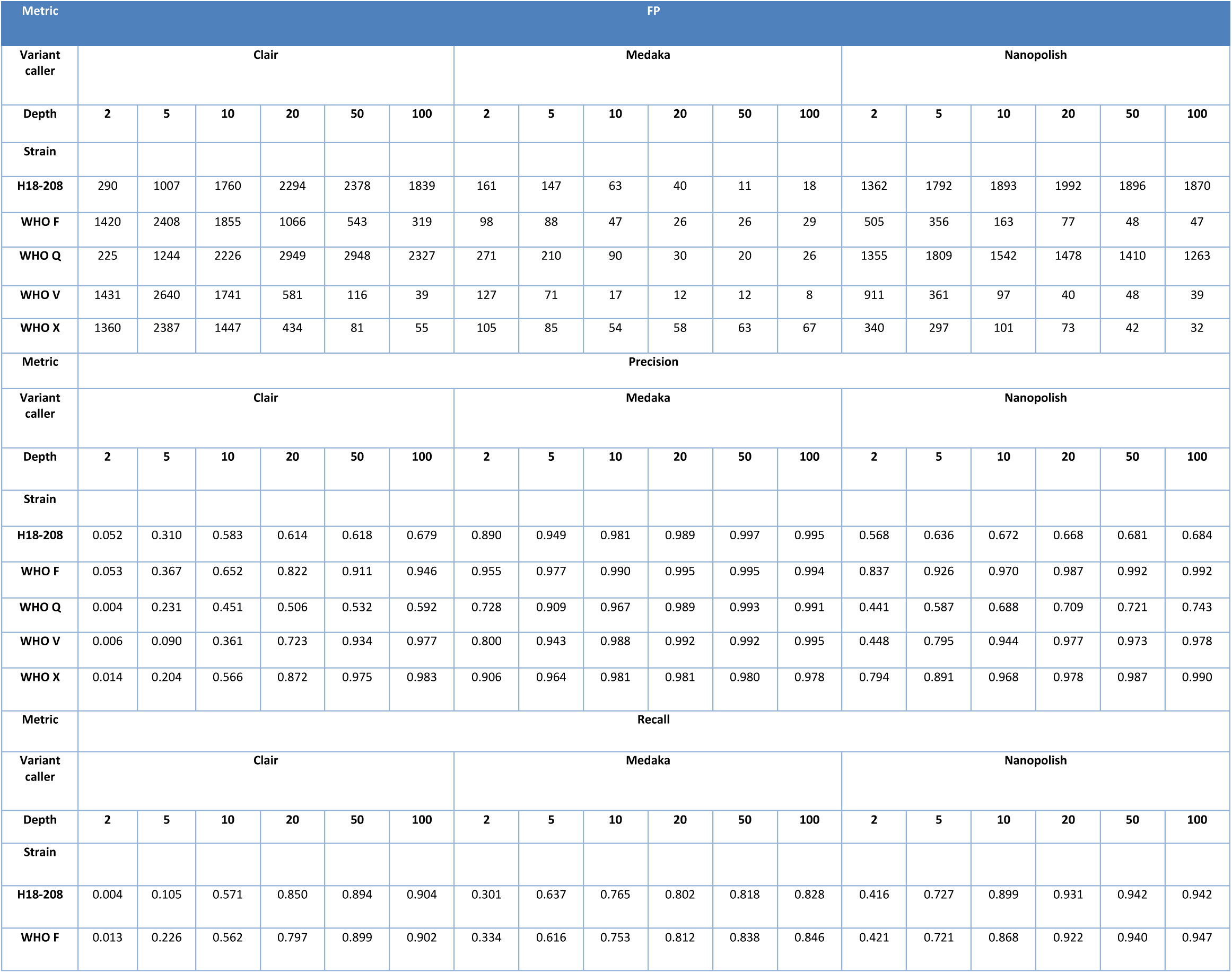
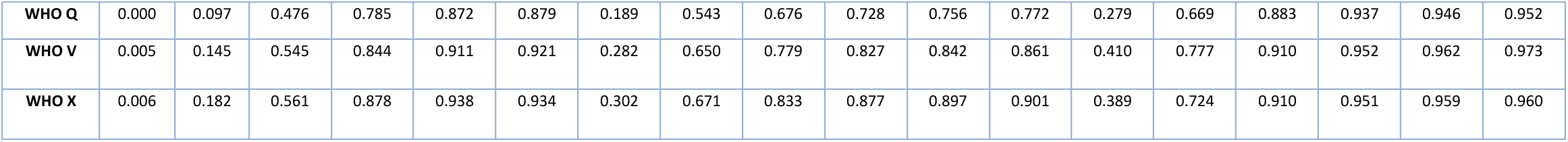
FP rate, Recall and Precision using a simple QUAL based threshold for filtering variant calls.

**Supplemental Table 3.** Discreet A2059G mutations discovered in 23s rRNA variant calls. Strains without 23s rRNA mutations are not shown. The expected number of A2059G mutations in WHO Q and WHO V is 4, as shown in bold.

| Strain | Depth | Discreet A2059G copies |  |  |
| --- | --- | --- | --- | --- |
|  |  | Nanopolish | Medaka | Clair |
| WHO Q | 2 | 2 | 2 | 2 |
|  | 5 | 3 | 3 | **4** |
|  | 10 | **4** | 3 | **4** |
|  | 20 | **4** | **4** | **4** |
|  | 50 | **4** | **4** | **4** |
|  | 100 | **4** | **4** | **4** |
| WHO V | 2 | 0 | 1 | 2 |
|  | 5 | 3 | 2 | **4** |
|  | 10 | **4** | **4** | **4** |
|  | 20 | **4** | **4** | **4** |
|  | 50 | **4** | **4** | **4** |
|  | 100 | **4** | **4** | **4** |

**Supplemental Table 4.** Genes and expected mutations for each strain at varying subsampled depths using Nanopolish or medaka for variant calling. Residues highlighted in yellow are incorrect.

| gene | Strain | Pos | Exp. Res. | Medaka |  |  |  |  |  | Nanopolish |  |  |  |  |  | Clair |  |  |  |  |  |
| --- | --- | --- | --- | --- | --- | --- | --- | --- | --- | --- | --- | --- | --- | --- | --- | --- | --- | --- | --- | --- | --- |
|  |  |  |  | 2 | 5 | 10 | 20 | 50 | 100 | 2 | 5 | 10 | 20 | 50 | 100 | 2 | 5 | 10 | 20 | 50 | 100 |
| gyrA | H18-208 | 91 | F | F | F | F | F | F | F | F | F | F | F | F | F | F | F | F | F | F | F |
|  |  | 95 | A | A | A | A | A | A | A | G | A | A | A | A | A | G | A | A | A | A | A |
|  | WHOF | 91 | S | F | F | S | S | S | S | F | F | S | S | S | S | F | F | S | S | S | S |
|  |  | 95 | D | D | D | D | D | D | D | G | D | D | D | D | D | G | G | D | D | D | D |
|  | WHOQ | 91 | F | F | F | F | F | F | F | F | F | F | F | F | F | F | F | F | F | F | F |
|  |  | 95 | A | G | A | A | A | A | A | G | A | A | A | A | A | G | A | A | A | A | A |
|  | WHOV | 91 | F | F | F | F | F | F | F | F | F | F | F | F | F | F | F | F | F | F | F |
|  |  | 95 | G | G | G | G | G | G | G | G | G | G | G | G | G | G | G | G | G | G | G |
|  | WHOX | 91 | F | F | F | F | F | F | F | F | F | F | F | F | F | F | F | F | F | F | F |
|  |  | 95 | N | G | G | G | G | G | G | G | N | N | N | N | N | G | N | N | N | N | N |
| parC | H18-208 | 86 | D | D | D | D | D | D | D | D | D | D | D | D | D | D | D | D | D | D | D |
|  |  | 87 | R | R | R | R | R | R | R | R | R | R | R | R | R | R | R | R | R | R | R |
|  |  | 88 | S | S | S | S | S | S | S | S | S | S | S | S | S | S | S | S | S | S | S |
|  | WHOF | 86 | D | D | D | D | D | D | D | D | D | D | D | D | D | D | D | D | D | D | D |
|  |  | 87 | S | S | S | S | S | S | S | S | S | S | S | S | S | S | S | S | S | S | S |
|  |  | 88 | S | S | S | S | S | S | S | S | S | S | S | S | S | S | S | S | S | S | S |
|  | WHOQ | 86 | D | D | D | D | D | D | D | D | D | D | D | D | D | D | D | D | D | D | D |
|  |  | 87 | R | R | R | R | R | R | R | R | R | R | R | R | R | R | R | R | R | R | R |
|  |  | 88 | S | S | S | S | S | S | S | S | S | S | S | S | S | S | S | S | S | S | S |
|  | WHOV | 86 | D | D | D | D | D | D | D | D | D | D | D | D | D | D | D | D | D | D | D |
|  |  | 87 | R | R | R | R | R | R | R | R | R | R | R | R | R | R | R | R | R | R | R |
|  |  | 88 | S | S | S | S | S | S | S | S | S | S | S | S | S | S | S | S | S | S | S |
|  | WHOX | 86 | D | D | D | D | D | D | D | D | D | D | D | D | D | D | D | D | D | D | D |
|  |  | 87 | R | R | R | R | R | R | R | R | R | R | R | R | R | R | R | R | R | R | R |
|  |  | 88 | P | P | P | P | P | P | P | P | P | P | P | P | P | P | P | P | P | P | P |
| ponA | H18-208 | 421 | P | P | P | P | P | P | P | P | P | P | P | P | P | P | P | P | P | P | P |
|  | WHOF | 421 | L | L | L | L | L | L | L | L | L | L | L | L | L | L | L | L | L | L | L |
|  | WHOQ | 421 | P | P | P | P | P | P | P | P | P | P | P | P | P | P | P | P | P | P | P |
|  | WHOV | 421 | P | P | P | P | P | P | P | P | P | P | P | P | P | P | P | P | P | P | P |
|  | WHOX | 421 | P | P | P | P | P | P | P | P | P | P | P | P | P | P | P | P | P | P | P |
| mtrR | H18-208 | 45 | G | G | G | G | G | G | G | G | G | G | G | G | G | G | G | G | G | G | G |
|  | WHOF | 45 | G | G | G | G | G | G | G | G | G | G | G | G | G | G | G | G | G | G | G |
|  | WHOQ | 45 | D | G | D | D | D | D | D | D | D | D | D | D | D | D | D | D | D | D | D |
|  | WHOV | 45 | G | G | G | G | G | G | G | G | G | G | G | G | G | G | G | G | G | G | G |
|  | WHOX | 45 | G | G | G | G | G | G | G | G | G | G | G | G | G | G | G | G | G | G | G |
| rpsJ | H18-208 | 57 | M | M | M | M | M | M | M | M | M | M | M | M | M | M | M | M | M | M | M |
|  | WHOF | 57 | V | V | V | V | V | V | V | V | V | V | V | V | V | V | V | V | V | V | V |
|  | WHOQ | 57 | M | M | M | M | M | M | M | M | M | M | M | M | M | M | M | M | M | M | M |
|  | WHOV | 57 | M | M | M | M | M | M | M | M | M | M | M | M | M | M | M | M | M | M | M |
|  | WHOX | 57 | M | M | M | M | M | M | M | M | M | M | M | M | M | M | M | M | M | M | M |

**Supplemental Table 5.**
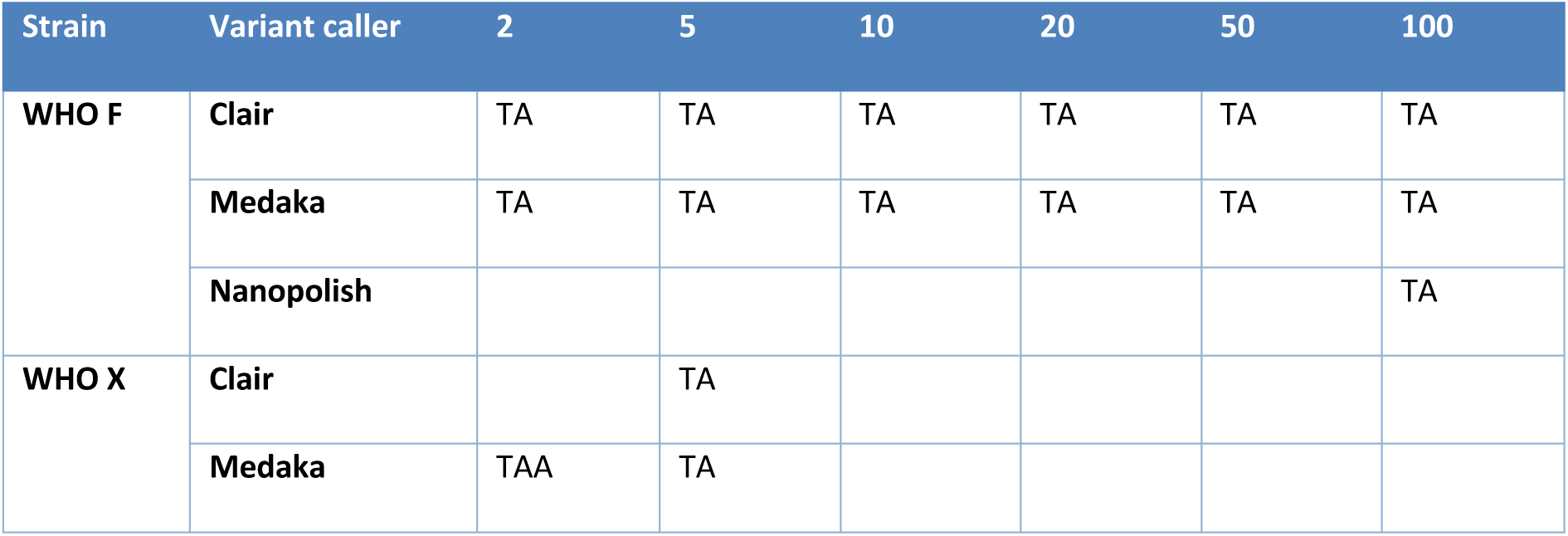
mtrR promoter sequences. ALT sequences identified at positions 1332810 in reference NC_011035.1 by each variant caller at different sub sampled depths.

| Strain | Variant caller | 2 | 5 | 10 | 20 | 50 | 100 |
| --- | --- | --- | --- | --- | --- | --- | --- |
| WHO F | Clair | TA | TA | TA | TA | TA | TA |
|  | Medaka | TA | TA | TA | TA | TA | TA |
|  | Nanopolish |  |  |  |  |  | TA |
| WHO X | Clair |  | TA |  |  |  |  |
|  | Medaka | TAA | TA |  |  |  |  |

**Supplemental Table 6.** Detection of penA using de novo assemblies. penA alleles were determined from each strain (Strain), detected from two different assembly approaches (Method), over a range of 5 different sub sampled depths (Depth N). Whole genome assembly (WGA) was performed with RA and local assemblies (LA) were performed with WTBDG2. Where identified the penA was correct.

| Strain | Method | 2 | 5 | 10 | 20 | 50 | 100 |
| --- | --- | --- | --- | --- | --- | --- | --- |
| H18-208 | WGA |  |  | 60.001 | 60.001 | 60.001 | 60.001 |
|  | LA |  |  | 60.001 | 60.001 | 60.001 | 60.001 |
| WHO F | WGA |  |  |  |  | 15.001 |  |
|  | LA |  | 15.001 | 15.001 | 15.001 | 15.001 | 15.001 |
| WHO Q | WGA |  |  |  | 60.001 | 60.001 | 60.001 |
|  | LA |  | 60.001 | 60.001 | 60.001 | 60.001 | 60.001 |
| WHO V | WGA |  |  | 5.002 | 5.002 | 5.002 | 5.002 |
|  | LA |  | 5.002 | 5.002 | 5.002 | 5.002 | 5.002 |
| WHO X | WGA |  |  | 37.001 | 37.001 | 37.001 | 37.001 |
|  | LA |  | 37.001 | 37.001 | 37.001 | 37.001 | 37.001 |

**Supplemental Table 7.**
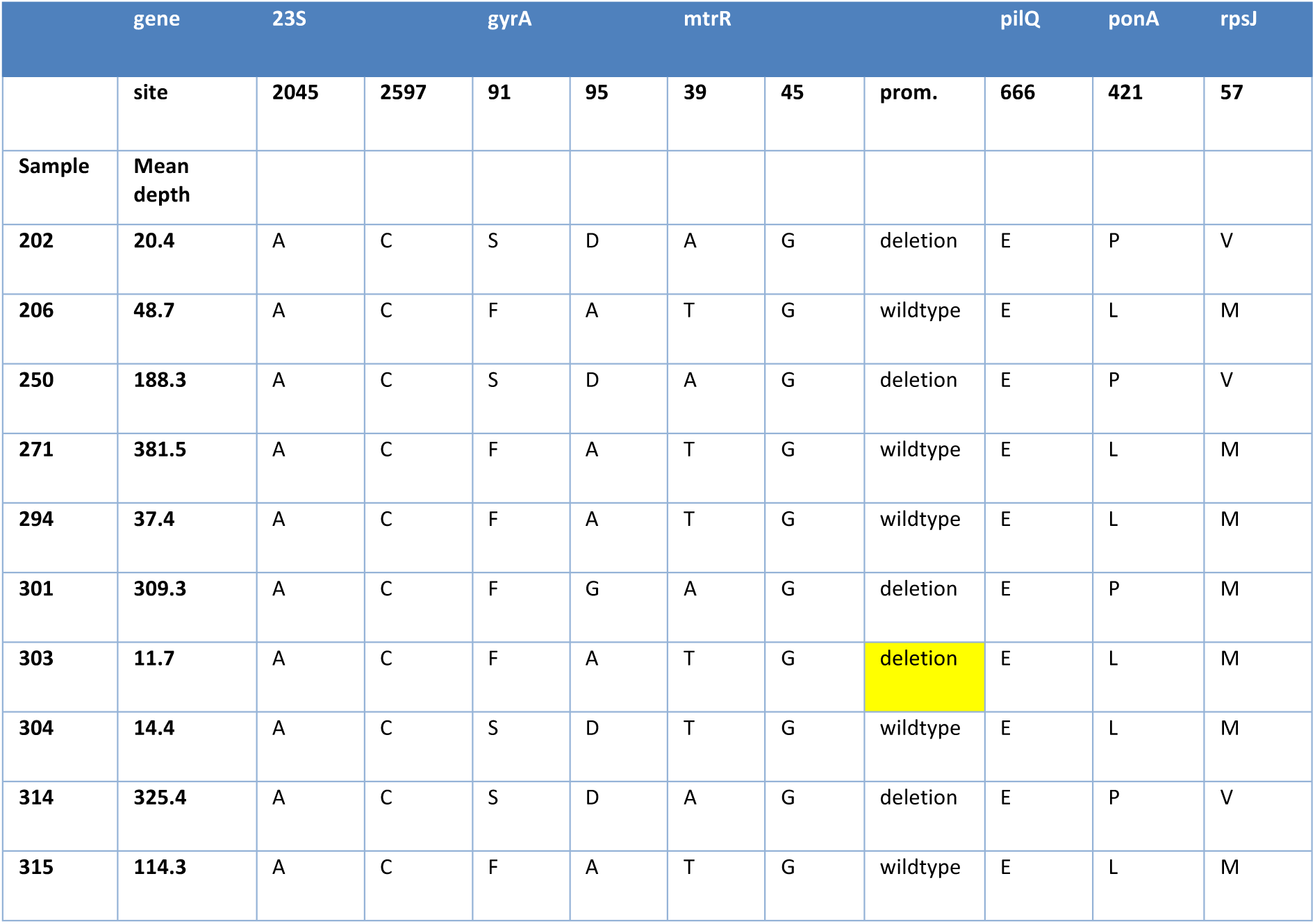
Antimicrobial resistance determinant detection in clinical samples. Clinical sample names, average coverage depth over the entire genome and the nucleotide base or amino acid residue detected for several important AMR genes. Yellow highlights where the nanopore results were not consistent with the Illumina results.

**Supplemental Table 8.** penA alleles identified by nanopore from clinical metagenomic samples and validated with Illumina sequenced cultures.

| Sample name | Allele | AMR Markers | Beta-Lactamase |
| --- | --- | --- | --- |
| 202 | penA2.001 | penA Type II NonMosaic | Negative |
| 206 | penA43.002 | penA Type 43 NonMosaic; A502V | Positive |
| 250 | penA2.001 | penA Type II NonMosaic | Negative |
| 271 | penA43.002 | penA Type 43 NonMosaic; A502V | Positive |
| 294 | penA43.002 | penA Type 43 NonMosaic; A502V | Positive |
| 301 | penA5.002 | penA Type V NonMosaic | Negative |
| 304 | penA14.001 | penA Type XIV NonMosaic | Negative |
| 314 | penA2.001 | penA Type II NonMosaic | Negative |
| 315 | penA5.002 | penA Type V NonMosaic | Negative |

